# Fast and interpretable scRNA-seq data analysis

**DOI:** 10.1101/2020.10.05.314039

**Authors:** Murat Can Çobanoğlu

## Abstract

One of the key challenges in single-cell data analysis is the annotation of cells with their cell types. This task is divided into two different sub-tasks: identifying known cell types and identifying novel cell types. In the former case, we can benefit from being able to transfer annotations from bulk RNA-seq because there are many more types profiled with that more established technology. In the latter case, we would benefit from interpretable models that can describe the reasons for grouping a number of cells together. We propose that both of these problems can be solved by generative Bayesian Dirichlet-multinomial models. In the supervised learning context, we propose a generative Bayesian Dirichlet-multinomial classifier. We show that such a classifier can effectively transfer cell labels from bulk to single-cell RNA-sequencing data. We also show that alternative well-established machine learning models have difficulty with this transition, even if they are effective within the same regime (i.e. single cell to single cell). In the unsupervised learning context, we propose a Bayesian Dirichlet-multinomial mixture model. We show that the proposed model learns meaningful clusters where the automatically learned relationships between cell types and genes overlap with ground truth associations. Furthermore, there are no density or connectivity based clustering assumptions in this model, which differs with almost every approach in this field. Consequently the clustering results from the generative method can effectively represent nuanced differences among cells.

## 1 Introduction

The advent of single cell RNA-sequencing (scRNA-seq) technology has enabled deeper understanding in a wide range of biomedical disciplines such as neuro-science (Welch *et al*., 2019), immunology (Jerby-Arnon *et al*., 2018), or oncology (van Galen *et al*., 2019). The ability to make observations at the single cell resolution offers unique advantages that are particularly useful in contexts comprised of complex multicellular architectures. Therefore the utilization of scRNA-seq has increased rapidly and is likely to increase further in the future.

However, scRNA-seq also offers unique challenges such as the necessity to annotate individual cells, low read counts per cell, or high number of data points (samples in bulk, cells in scRNAseq). Oftentimes, the very advantages themselves lead to challenges. For instance, observing transcriptomic phenotypes at the single cell resolution offers the ability to dissect cellular differences in heterogeneous contexts. However this necessitates the attribution of cell identity to each cell as an added step in data analysis. Observing a large number of cells is advantageous, yet this also demands efficient algorithms that can scale efficiently. Finally, one of the main challenges in the field is the development of interpretable machine learning methods that can communicate the learned biological insights to the user.

With current techniques, the users often have to make a choice between efficiency and interpretability. Previous Bayesian generative models such as BISCUIT (Prabhakaran *et al*., 2016) or scVI (Lopez *et al*., 2018) offer interpretable models with easily understandable insights. However the approximate inference methods used in these methods (variational inference in scVI, Gibbs sampling in BISCUIT) require at least hours of computation time for just one thousand cells. Biomedical scientists can now routinely collect data from tens of thousands of cells, with new initiatives such as the Human Cell Atlas planning to collect 30 million to 100 million (Regev *et al*., 2018). Therefore efficient alternatives are required to scale to large cell counts.

Due to the large numbers of cells, users of scRNA-seq often have to rely on scalable approaches. Consequently, despite the presence of hundreds of alternative scRNA-seq clustering methods practioners have previously used standard clustering methods such as k-means (Burns *et al*., 2015; Guo *et al*., 2017) or DBScan (Tirosh *et al*., 2016). The popular Seurat package (Butler *et al*., 2018; Stuart *et al*., 2019) utilizes connectivity-based clustering similar to PhenoGraph (Levine *et al*., 2015)and SNN-Cliq (Xu and Su, 2015). While useful and relatively efficient, these approaches have two main drawbacks compared to generative Bayesian approaches. The first is that they offer little generative insight that generative methods can, such as which specific genes led to the co-clustering of a number of cells. Secondly, they do not allow for natural prior incorporation in the manner of a Bayesian model.

The lack of priors can present subtle challenges. For instance, the presence of a signal spread over hundreds of genes (ex: metabolic genes) can overwhelm expression from a smaller set of genes (ex: cluster of differentiation genes) even if these handful are *a priori* known to be more informative. To illustrate, consider two T cells and two B cells such that one cell of each type is hypoxic while the other is normoxic. An immunologist manually annotating cell types would ignore the metabolic genes and consider the CD3 and CD19 expression of these cells. However, the strong effect of metabolic differences can easily skew a neighborhood-based or global similarity based approach. An interpretable Bayesian approach, in contrast, would enable the user to input the prior information to appropriately inform the method. Furthermore, an interpretable Bayesian method would, even under failure, be able to communicate the reasons for its decisions transparently by annotating the genes that define each cell type. Due to these considerations, we posit that scalable Bayesian models are necessary in this domain.

Finally, we argue that the methods for analyzing new single-cell RNA-sequencing methods should be able to utilize the wealth of existing bulk RNA-seq studies. There is pioneering work in this direction (Unified RNA-Seq Model; URSM) due to Zhu and colleagues (Zhu *et al*., 2018) where they use bulk to impute dropouts in single cell data, and in turn use single cell to deconvolve bulk samples. However they consider the cell labels in single cell to be known and indeed require its input to the model, therefore this approach is not suitable to the task of cell type annotation that we focus on. The vast majority of the literature that exists on single cell RNA-seq data analysis focuses exclusively on single cell data.

We hereby present two computational models (supervised and unsupervised) for fast generative Bayesian cell type annotation in scRNA-seq data (Figure 1). Our formulation rests on an overdispersed Dirichlet-multinomial generative discrete distribution in both cases. Briefly, the driving intuition is to propose an interpretable yet expressive generative formulation and then mathematically derive computationally efficient learning operations. We propose a supervised learning model (classifier) that enables cell identities to be learned from bulk RNA-seq and transferred to single cell RNA-seq. We then propose an unsupervised learning model (clustering) that enables the grouping of cells in scRNA-seq data.

**Figure 1:**
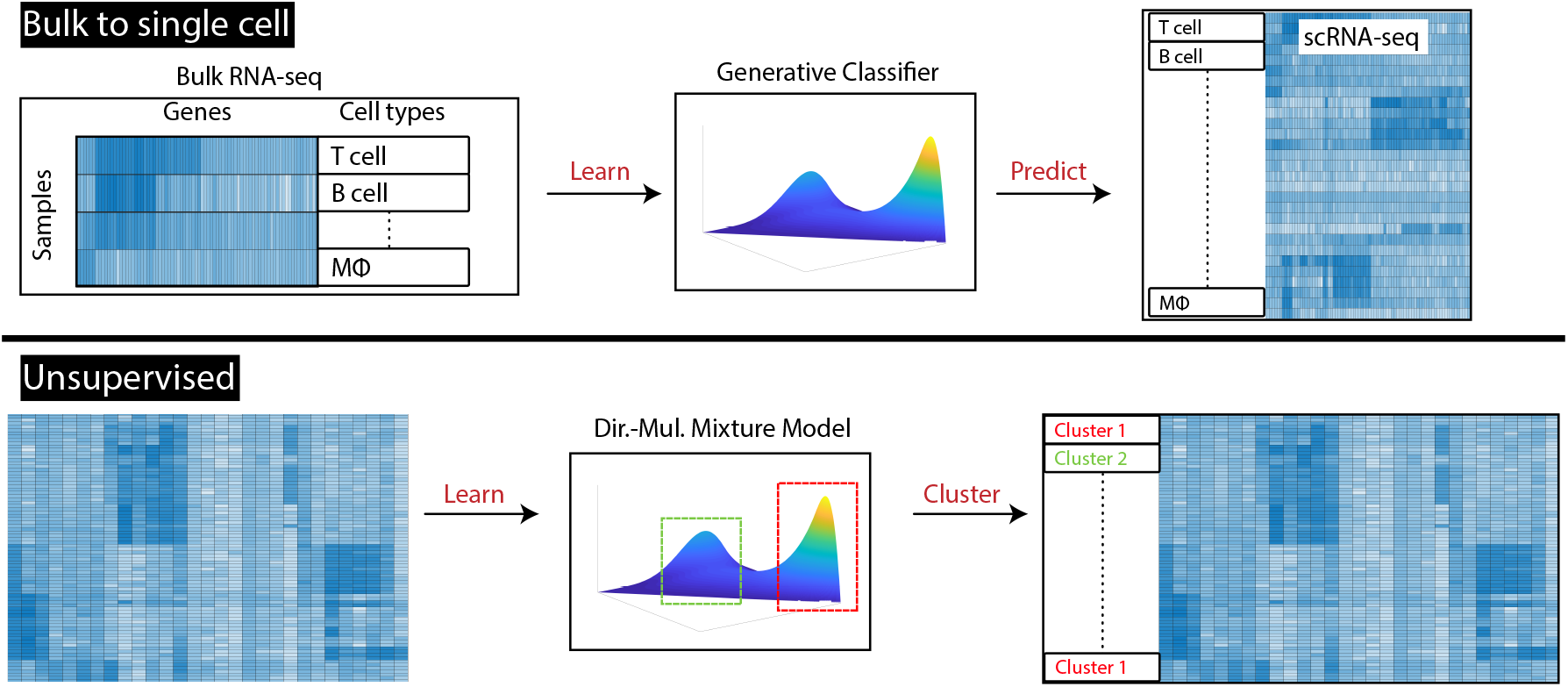
Overview of the proposed approach. In the supervised context, we learn a generative Dirichlet-multinomial model, potentially from bulk transcriptomic data, and then apply that to single-cell data. This is useful when seeking to assign *a priori* known cell type labels. In the unsupervised regime, we learn a Dirichlet-multinomial mixture model for clustering cells. This informs cluster labels to groups of cells, which can identify similarities/differences across a handful of genes if necessary, and also explain the decision.

We envision that practitioners who collect new scRNA-seq data would apply these two models in sequence. To elaborate, the user would first make the cell type calls that can be made with low uncertainty. Here they would use a major benefit of our method’s probabilistic model, which is that it natively supports uncertainty calculations. Then, the practitioner would use unsupervised learning to cluster the remaining uncalled cells into similar groups to detect potentially novel cell types. However our formulation is fully flexible and permits alternative uses such as label transfer (i.e. classification) alone or clustering alone. In the following sections, we both detail our method and describe the results that show that we have achieved both of these objectives.

## 2 Methods

Our approach centers on fast Newton-Raphson (NR) optimization for efficiently learning the parameters of a Dirichlet-multinomial (Pólya) distribution. In section 2.2, we discuss the mathematical details of this fast inference protocol as well as how we learn a class-conditional Dir.-mul. model for classification. In section 9(a), we extend this formulation to the mixture model setting, which we solve using the Expectation-Maximization algorithm. Jointly, we present methods for both supervised and unsupervised cell type label assignment for single-cell data.

### 2.1 Notation

We use the 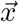 notation to represent that *x* is a vector. We denote the *i*-th element of the vector 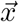with *x*^(*i*)^, while we index different vectors with subscripts such as 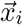 and 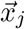. While the subscript is commonly chosen to represent the elements of a vector or matrix, we adopt this switched representation because we will more frequently need to index entire vectors (specifically, the concentration parameters of the Dirichlet distribution) than we will individual elements. We will use 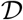 to represent the data and 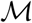 to represent all the parameters associated with the model. We will commonly represent constant factors that do not effect optimization (and thus model learning) with *C* without necessarily specifying all their constituents. These will often be related to the hyperparameter-only normalization terms for the various probability distributions we use for the priors. Finally, we will use *p*_*Exp*_ and *p*_*Dir*_ to denote, respectively, the PDF of the exponential and Dirichlet distributions while *p*_*DirMul*_ will denote the PMF of the Dirichlet-multinomial distribution.

### 2.2 Bayesian Dirichlet-multinomial generative classification

#### 2.2.1 Definition

We propose the following Bayesian generative model:

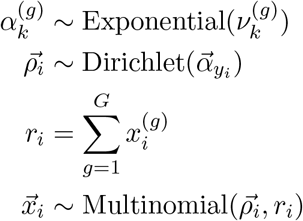

where 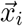 represents the read count vector over all the genes for the *i*-th data point (sample in bulk, cell in single cell), *y*_*i*_ represents the class label for the *i*-th data point, 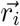 represents the total read count per sample, 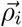 represents the probability vector such that the probability of sample *i* expressing gene *g* is 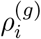, 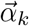 represents the vector of concentration parameters specifying the Dirichlet distribution that corresponds to the *k*-th class, and 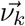 represents the hyperparameters for specifying prior belief for 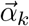 elementwise. In all the results in this paper, we set 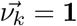 and thus utilize the Bayesian formulation for regularization alone. However, the users would be able to set it arbitrarily in order to deliver *a priori* available model insight, such as increasing the propensity for high concentration parameters for known marker genes of a given cell type. A graphical representation of the described model is provided in Figure 2.

**Figure 2:**
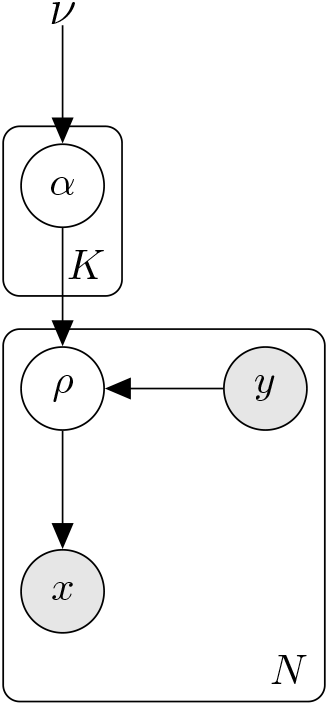
Graphical representation of the class-conditional generative classifier model.

#### 2.2.2 Inference

We will learn the *maximum a posteriori* (MAP) estimate of the model parameters using the Newton-Raphson (NR) optimization method. The key parameters to learn in the supervised case are the 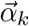 that parameterize the class-conditional Dirichlet distributions, specifying the gene expression propensities. To learn these parameters using NR, we need to compute the following update rule:

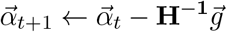

where **H**^−**1**^ and 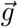 are, respectively, the inverse of the Hessian and the gradient of the objective function (the log-joint *l* in our case) which are defined as follows:

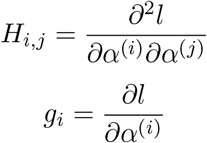

To derive the update rule, we can write the joint probability of the data and the parameters as:

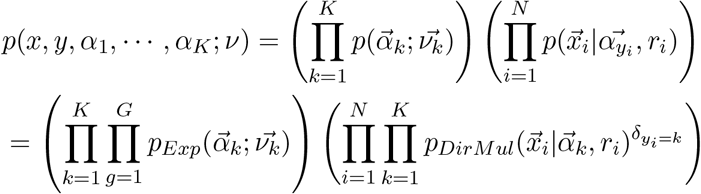

where *N* is the number of data points (cells or samples), *K* is the number of classes, *G* is the number of genes, *δ* is the Dirac delta function, and *p*_*Exp*_ and *p*_*DirMul*_ are respectively the probability distribution function (PDF) of the exponential distribution and the probability mass function (PMF) of the Dirichlet-multinomial compound distribution.

We then use the definitions of exponential PDF and the Dirichlet-multinomial PMF distributions, and apply the logarithmic transform to write the log-joint as:

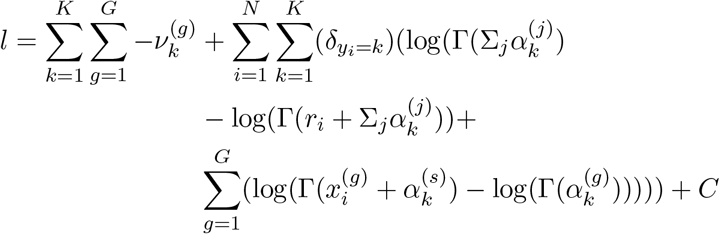

where 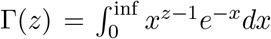 and *C* represents all the constants that do not vary with the model parameters. Since logarithm is a monotonic transformation the optima of the log-joint are equivalent to the joint and, as is almost universally the case, we will optimize the log-joint for numerical stability. To derive the update rules, we can then write the partial derivatives that constitute the Hessian and the gradient as:

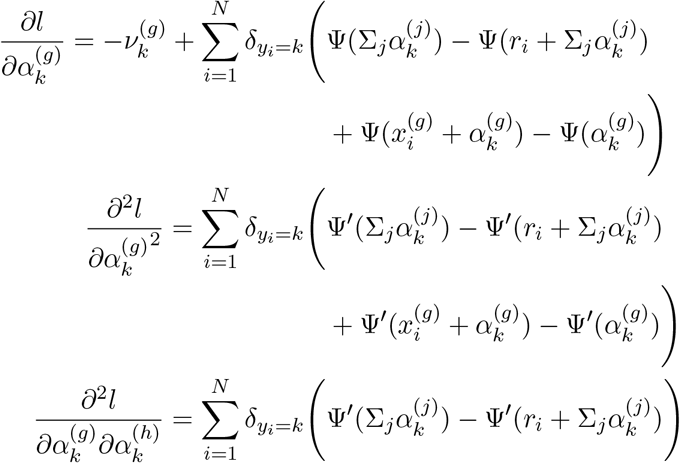

where Ψ and Ψ′ are respectively the digamma and the trigamma functions defined as 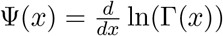 and 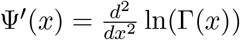. To efficiently optimize this function, we follow some of the ideas presented in (Minka, 2012), and observe that we can rewrite the Hessian **H** as follows:

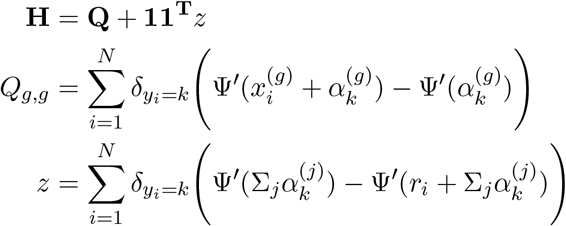

where **Q** is a diagonal matrix, *z* is a constant, and **1** is a vector of 1s of length *G*. This enables us to use the Sherman-Morrison formula to write:

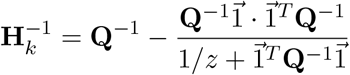

which enables us to calculate 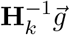 directly in linear time w.r.t. *G*. Our formulation enables us to use highly optimized vectorized implementations (TensorFlow, Matlab etc.) of all the fundamental mathematical operations necessary for computing the update rules. Furthermore having only linear time dependency on the number of genes enables this model to scale to a high number of genes. Therefore the above described learning process can be implemented to be highly efficient.

Finally, because we use NR instead of gradient ascent, each update step contributes significantly more information to the learning process. Therefore in practice substantially fewer steps are necessary for convergence. Another advantage of this learning procedure is that there are no model learning hyperparameters (such as “step size” or “momentum”). Therefore the user can simply use the inference procedure out-of-box, with no data or domain specific parameter tuning.

#### 2.2.3 Classification

Given a model 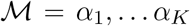, we can predict the class membership of any new data vector 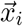 by simply identifying *k* that maximizes the class-conditional likelihood:

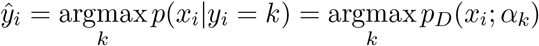

One of the main advantages of having a fully probabilistic Bayesian generative model is that for any data point we can interrogate its probability of membership to all alternative classes. This can have useful applications such as using the entropy (Σ_*k*_ −*p*_*k*_ log(*p*_*k*_)) of the label predictions to decide when to call cell types and when to declare them unclassified. When assigning cell types to cells in scRNA-seq, this is useful in determining which cells can be novel cell types and which can be confidently called as one of the *a priori* known classical cell types.

### 2.3 Bayesian Dirichlet-multinomial mixture model clustering

#### 2.3.1 Definition

In the unsupervised learning regime, we propose the following Bayesian Dirichlet-multinomial mixture model (BaD3M; pronounced *Bah-dem*) generative model:

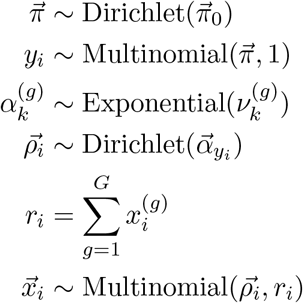

where the main difference with the supervised classification model in Section 2.2 is that we know consider the label of every data point, *y*_*i*_, to be a latent variable instead of an observed variable. In conjunction with this change, we introduce 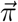 as a Dirichlet distributed latent random vector that represents the mixture weights for the various components. A graphical representation of the described model is provided in Figure 3.

**Figure 3:**
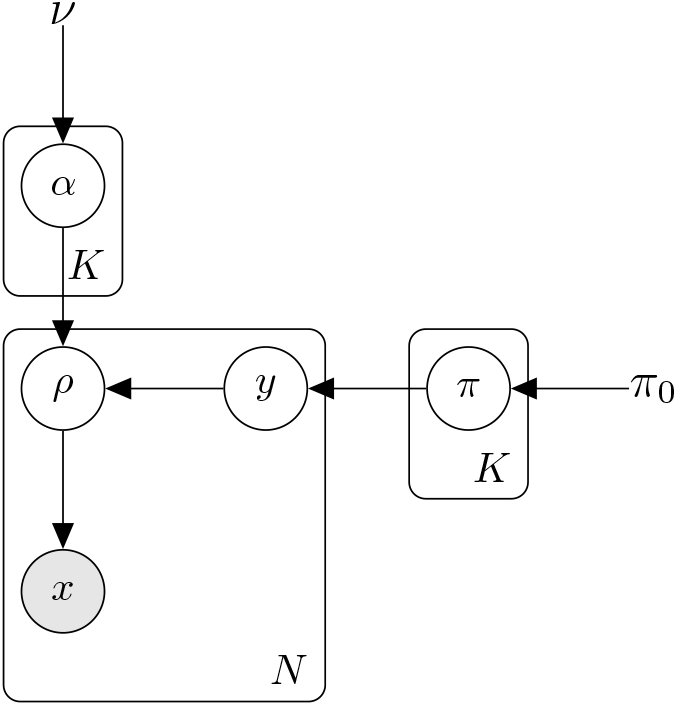
Graphical representation of the generative Dirichlet-multinomial mixture model.

#### 2.3.2 Inference

We use the Expectation-Maximization (EM) algorithm to learn the model parameters. Here, we write the joint distribution of the data 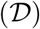 and the model parameters 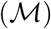 as:

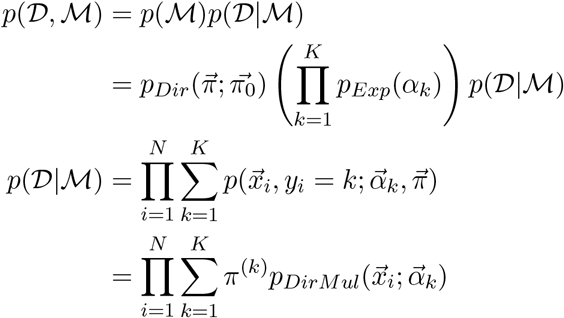

where the most important difference from the model in Section 2.2 is that we introduce *y*_*i*_ to be the latent variable that denotes the mixture component from which the *i*-th data point was generated (i.e. sample in bulk, cell in scRNA-seq) belongs. We will use this latent variable to assign clusters to the data points after we are done with the inference. Additionally, the mixture weights are written as 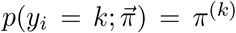. Given this expression, we can write a lower bound on the log of the joint distribution as:

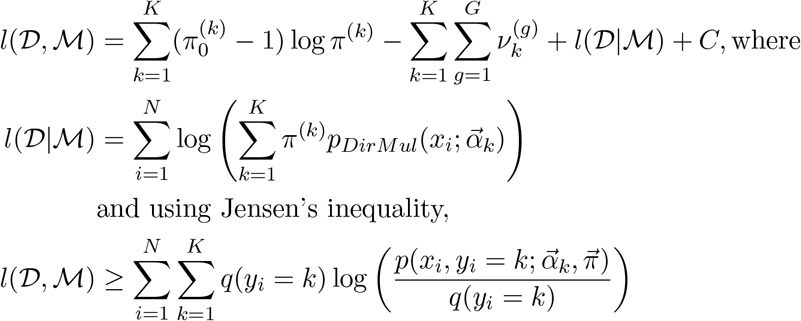

where we define *q*(*y*_*i*_ = *k*) = *p*(*y*_*i*_ = *k*|*x*_*i*_; *ϕ*) to be the class posterior probabilities with *ϕ* = {*π, α*_1_, … , *α*_*K*_} being the model parameters. In expanded form, we can then bound the log-joint as follows:

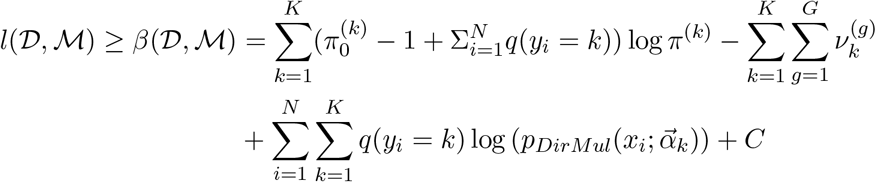

To learn the mixture weights *π*^(*k*)^ that maximize this lower bound, we need to set 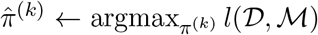 subject to the constraint that Σ_*k*_ *π*^(*k*)^ = 1. To solve this optimization problem, we construct the following Langrangian:

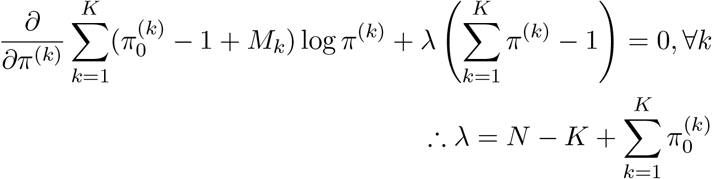

where 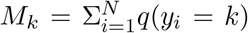. To learn the 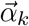, we can use a similar approach to that for the classifier. Specifically, we can derive the gradient and the constituents of the Hessian as:

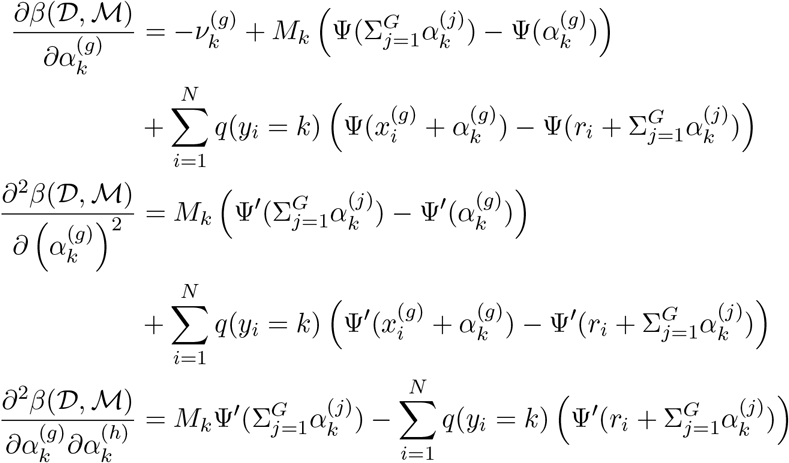

We can then iteratively optimize *α*_*k*_ using the Sherman-Morrison trick based Newton-Raphson updates as we did in Section 2.2.2. Specifically, we can now write:

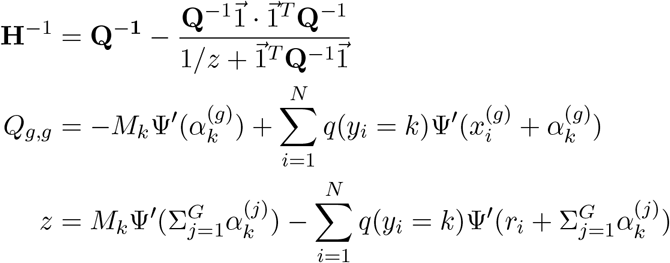

By integrating these ideas, we can then write our EM algorithm as:

<u>E-step</u>:

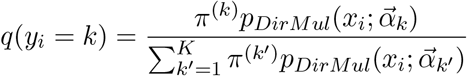

<u>M-step</u>:

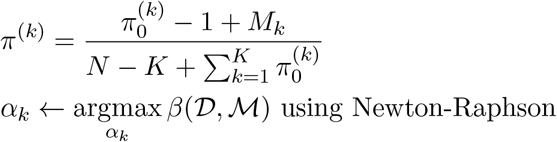

#### 2.3.3 Clustering

After learning the model, we can compute the posterior class probabilities *p*(*y*_*i*_ = *k*|*x*_*i*_) as follows:

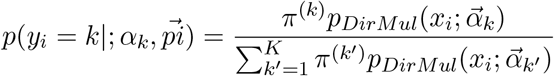

The posterior probabilities can be used to assign clusters to individual data points simply by choosing the maximum:

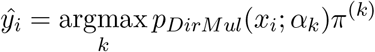

It should be noted that having a probabilistic cluster assignment can deliver much more utility than the cluster assignments. For instance, it is possible to compute the uncertainty in cluster assignments, or identify if a cell was comparably likely to be a member of two or more clusters.

## 3 Results

The models we present can be used for many different applications, either individually or in combination. To highlight their impacts on some of the most common tasks, we organized our results in two parts. First, we discuss the results when using the generative model to transfer cell type labels from bulk RNA-seq to scRNA-seq (section 3.1) whereas alternative methods cannot. This demonstrates the success with which the aforementioned model can cross these two significantly different regimes. This is also the first method that accomplishes this cross-domain transfer, to the best of our knowledge. We then take an unlabeled group of cells and show that our method can correctly pick out the cells in each group (section 3.2). Finally, the probabilistic

### 3.1 Bulk to single cell label transfer

Cancer microenvironment consists of a highly heterogeneous collection of cell types (Tabassum and Polyak, 2015). Therefore scRNA-seq is highly useful in this domain. Within the cells in the cancer microenvironment, the cells with the best defined cell types are the lymphocytes. While the different subclones among malignant cells are disputable, the distinction between lymphocytes (say, T cell versus B cell) are clearly established. Given these, we chose to focus on tumor infiltrating lymphocytes to test our proposed classification model.

#### 3.1.1 BDMC automatically learns marker genes

To interpret the biological relevance of the inferred models, we considered the inferred class-specific gene expression models on bulk immune cell transcriptomes to the ground truth marker associations . For training the model, we used the genes that the authors of the single cell study we use as test data used as markers for specific cell types (Tirosh *et al*., 2016). Specifically, they assigned a set of genes as markers to various cell types (for instance CD2, CD3D/E/G for T cells) and assigned a cell type label to a population of cells if most of them seemed to have high average expression of those markers. They used knowledge about celltype to marker gene associations, whereas in this study we made a weaker assumption that only a set of relevant genes, but without their assignment to individual cell types. We then considered whether if BDMC could learn the known ground truth associations between cell types and their known markers automatically. In addition to the markers for all the lymphocytes, which comprised all the cell types in our training set, to potentially confuse the model we also included some markers for cell types not in the training. Examples include the endothelial marker VWF or fibroblast markers COL1A1 and COL3A1.

**Table 1:** Confusion matrix for BDMC trained on bulk RNA-seq and tested on single cell RNA-seq. Prediction accuracy is 98.1%. Training data are 57 bulk samples; test data are 85,423 FACS-sorted single cells, with varying purity (Macrophage: 98%, T cells: 95% to 99%, NK: 92%, B cells: 100%)

|  | B cell | Macro. | Mono. | NK cell | Neutrophil | T cell |
| --- | --- | --- | --- | --- | --- | --- |
| B cell | **10082** | 0 | 0 | 0 | 0 | 3 |
| Mono. | 320 | 122 | **2129** | 15 | 11 | 15 |
| NK cell | 12 | 2 | 2 | **8078** | 1 | 290 |
| T cell | 240 | 11 | 16 | 525 | 5 | **63544** |

Initially, we considered three well established marker genes and cell types as shown in Figure 4. We observed that our model was able to accurately infer Dirichlet PDF parameters that matched with the known biological relationships between these genes and cell types. We chose three genes because the PDF they define over a 2-simplex is easily visualizable.

**Figure 4:**
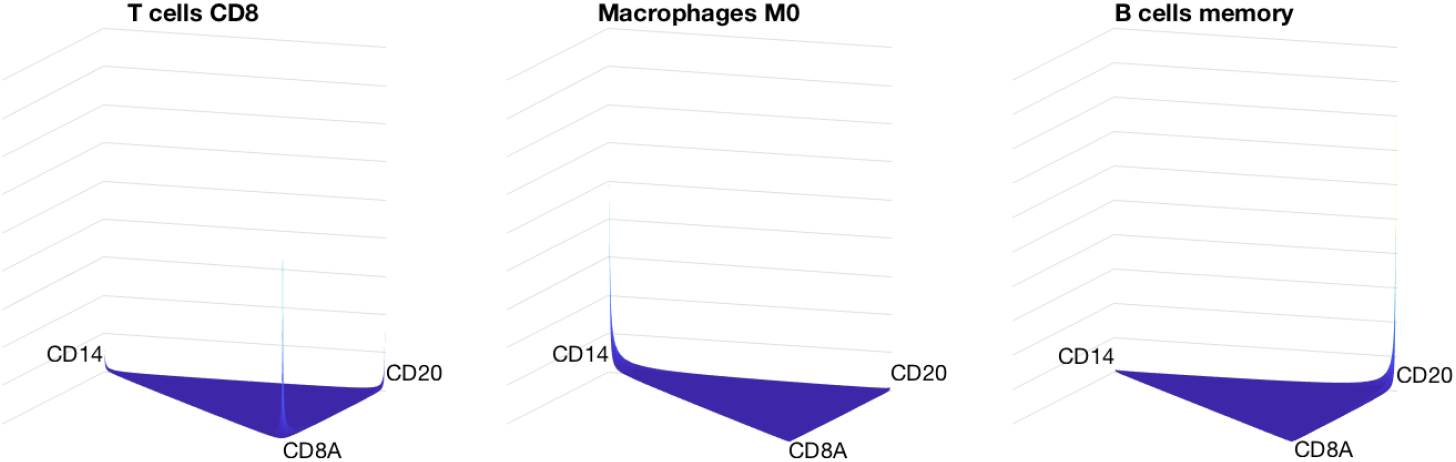
Inferred Dirichlet probability density function parameters reflect known biology. We show the probability density functions of tri-variate Dirichlet distributions using only the concentration weights learned for three marker genes for three specific cell types. CD8A, CD14, and CD20 are known marker genes for, respectively, CD8^+^ T cells, macrophages, and B cells. The model appropriately inferred the specific propensities for expressing each of these genes in each cell type.

Next, we considered all the markers for all the genes as shown in Figure 5. We observed that the model correctly matched each cell type with the marker genes known to belong to that cell type. For instance, pan T cell markers CD2 or CD3 were correctly associated with both CD8^+^ T cells and CD4^+^ T cells, while CD8A/B were correctly associated with only CD8^+^ T cells. The markers associated with cell types not included in the training set (such as collagen genes) were correctly learned to not be pertinent to any of the cell types.

**Figure 5:**
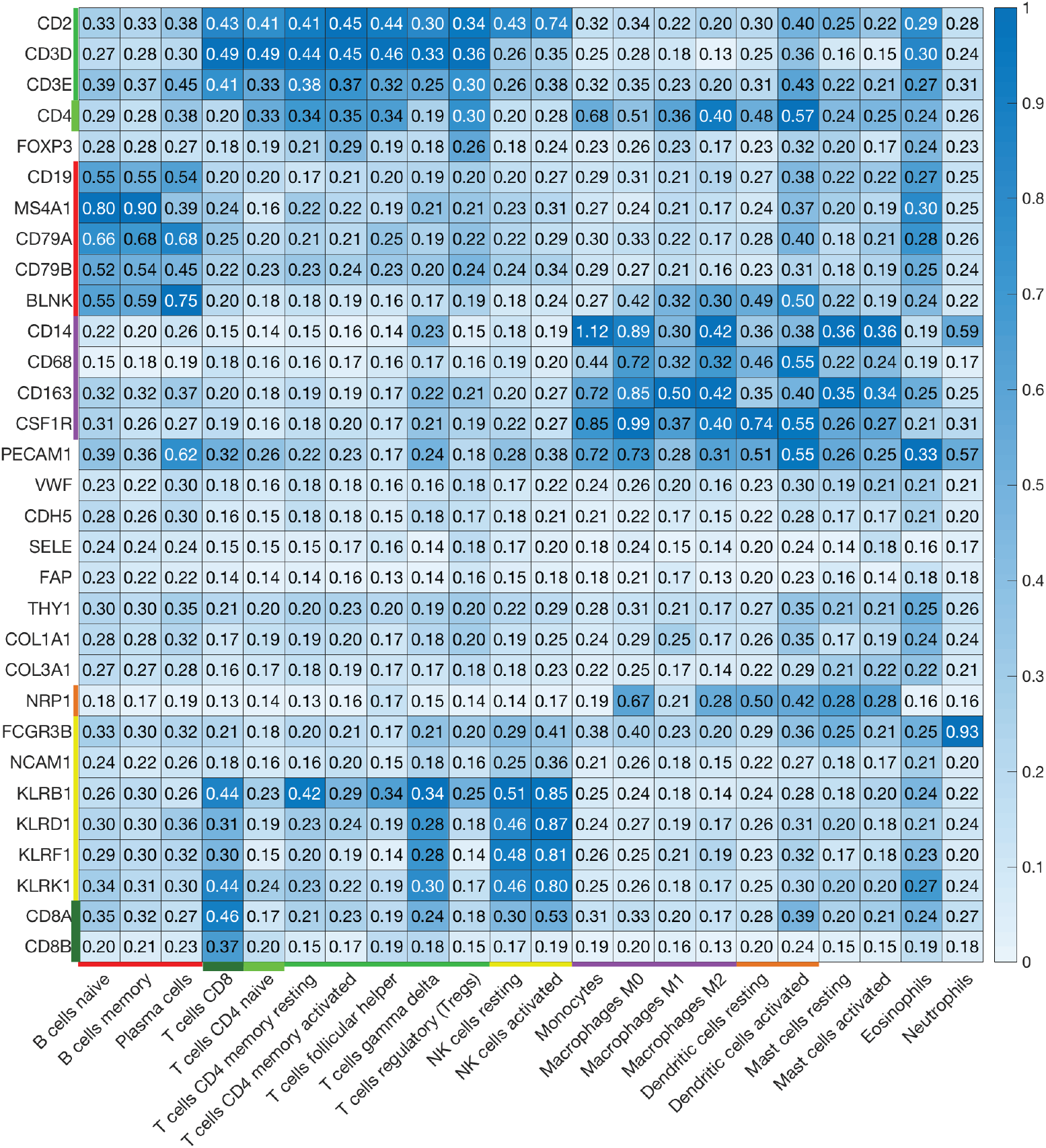
The latent variables that the BDMC model learns for the lympho-cyte population. To assess the quality of the learned model, we plot the 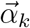 values for each class *k* after training on a set of bulk transcriptomic profiles of immune cell populations. The markers of specific cell types are indicated in the rows and columns with identically colored bars.

#### 3.1.2 BDMC transfers cell type information from bulk to single cell

Next, we hypothesized that the probabilistic formulation we used could accurately transfer cell type label information from bulk to single cell. The motivation behind this hypothesis was that our model learns an overdispersed, relative probabilistic model of each cell type completely dissociated from the read counts. Therefore, so long as the relative expression of genes were mostly preserved (a sensible assumption when considering cells of the same type) then the probabilistic model should hold equally well in bulk or single cell.

To train the model, we used the bulk transcriptomes of lymphocytes dataset curated by Newman and colleagues (2015) from numerous other studies in the literature. We would like to note here that not all of these samples are bulk RNA-seq *per se*, but also contain some microarray data. To demonstrate the strength of our approach in handling heterogeneous data inputs, we cast these transcriptomic profiles into integers to yield pseudo read counts.

To test the model, we used the melanoma tumor microenvironment scRNA-seq data collected by Tirosh and colleagues (2016). The authors already assigned labels through a combination of manual annotation based on markers and DBSCAN. We took these author assignments to be the ground truth and used them to assess the performance of each model.

Finally, we would like to note that there are some cell types in the training set that do not map to any of the labels in the single cell data (such as eosinophils, mast cells, etc). We deliberately left those cell types in the training dataset because when these methods are deployed on real world data, the users will not know exactly which cell types to expect in the single cell data. Therefore it is highly likely that they will train a model on more cell types than those that exist in the classification target dataset. Hence, a good model should be robust in avoiding these classification labels.

We compared our method against different alternatives that are commonly and successfully used in supervised learning, namely, decision forests (DF), support vector machines (SVM), and Gaussian discriminant analysis (GDA). Decision forests and support vector machines are flexible in dealing with a wide variety of supervised learning problems. We included GDA because it is an alternative generative method that uses a different formalism to represent the data.

BDMC was able to accurately transfer cell type information from bulk RNA-seq to scRNA-seq whereas other approaches could not (Figure 6). The zero accuracy stems from the fact that these models classified the single cells into classes that existed in the training data but not in the test data. To make sure that these results were not simply due to faulty parameterization, we used MATLAB implementations of these methods and used automatic hyperparameter optimization where applicable. The discrepancy between the read counts in the bulk and single cell regimes can dramatically reduce the performance of these methods. To eliminate facile errors due to this discrepancy, we used quantile normalization for these methods yet the results did not change. We present that our method, BDMC, handles this shift naturally since it receives read counts per sample (or per cell) as an input to the generative probability distribution and automatically scales.

**Figure 6:**
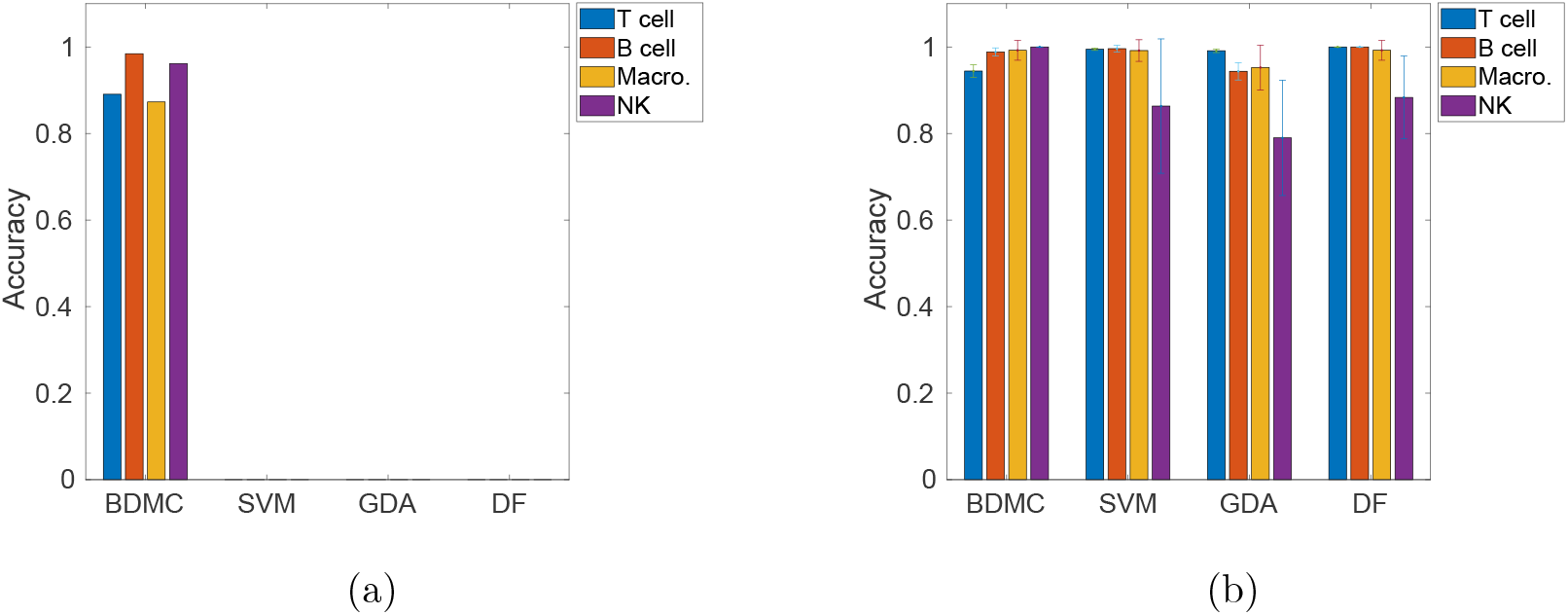
Classifier accuracy across and within bulk & single cell regimes. (a) We show accuracy of models trained on bulk and predicting single cell labels. BDMC can correctly transfer information from bulk to single cell, but other methods have difficulty. (b) We show the accuracy of training and testing a model on single cell data alone using k-fold cross validation (k=10; bars show average, error bars show standard deviation).

### 3.2 Single cell RNA-seq clustering

One of the most important motivations in collecting single cell RNA-sequencing data is to identify novel populations of cells that could not be detected before. Viewed from this lens, classification is useful in filtering out those cells that can be confidently called as one of the *a priori* known cell types. However, we still need a method for understanding similarities among those cells that cannot be called. In this context, we cannot expect to rely on labels therefore we need to perform unsupervised learning.

To estimate the performance of BaD3M in clustering unlabeled cells, we consider the tumor infiltrating lymphocytes without their labels and compare the clustering results to the ground truth labels. We restricted the genes mostly to the markers that the authors used for *a priori* class-specific assignment. We compare the author assigned labels to those acquired by our clustering method, as well as the results from the commonly used K means algorithm. At this point, we would like to note that the author assigned labels are not necessarily golden standard ground truth annotations. The authors grouped cells together and they assigned labels to all the cells in the same group based on the dominant markers, despite different expression profiles. While we expect most of the vast majority of their annotations to be accurate, there is some room for errors. Therefore we expect to see overall agreement but not necessarily perfect overlap.

To compare the results of various approaches at a single glance, we used the popular t-SNE (Maaten and Hinton, 2008) method to reduce dimensionality and visualize the data (Figure 7). We colored the cells with the clusters assigned by our method and k-means, in addition to showing their author-assigned labels. Our method was able to infer a clustering that, overall, matched quite well with the ground truth labels, where K means failed to do so. The dataset clearly consists of highly imbalanced clusters. Our proposed BaD3M method can handle this because the frequency of each class is a latent variable that we learn, through the *π* mixture weight parameters in the model. Furthermore, our method can learn that two clusters are very similar to each other on most genes, except for a large difference on a single gene which is the discriminative feature. Both of these are challenges for K means since it assumes uniformly sized clusters, and it has a density based clustering assumption.

**Figure 7:**
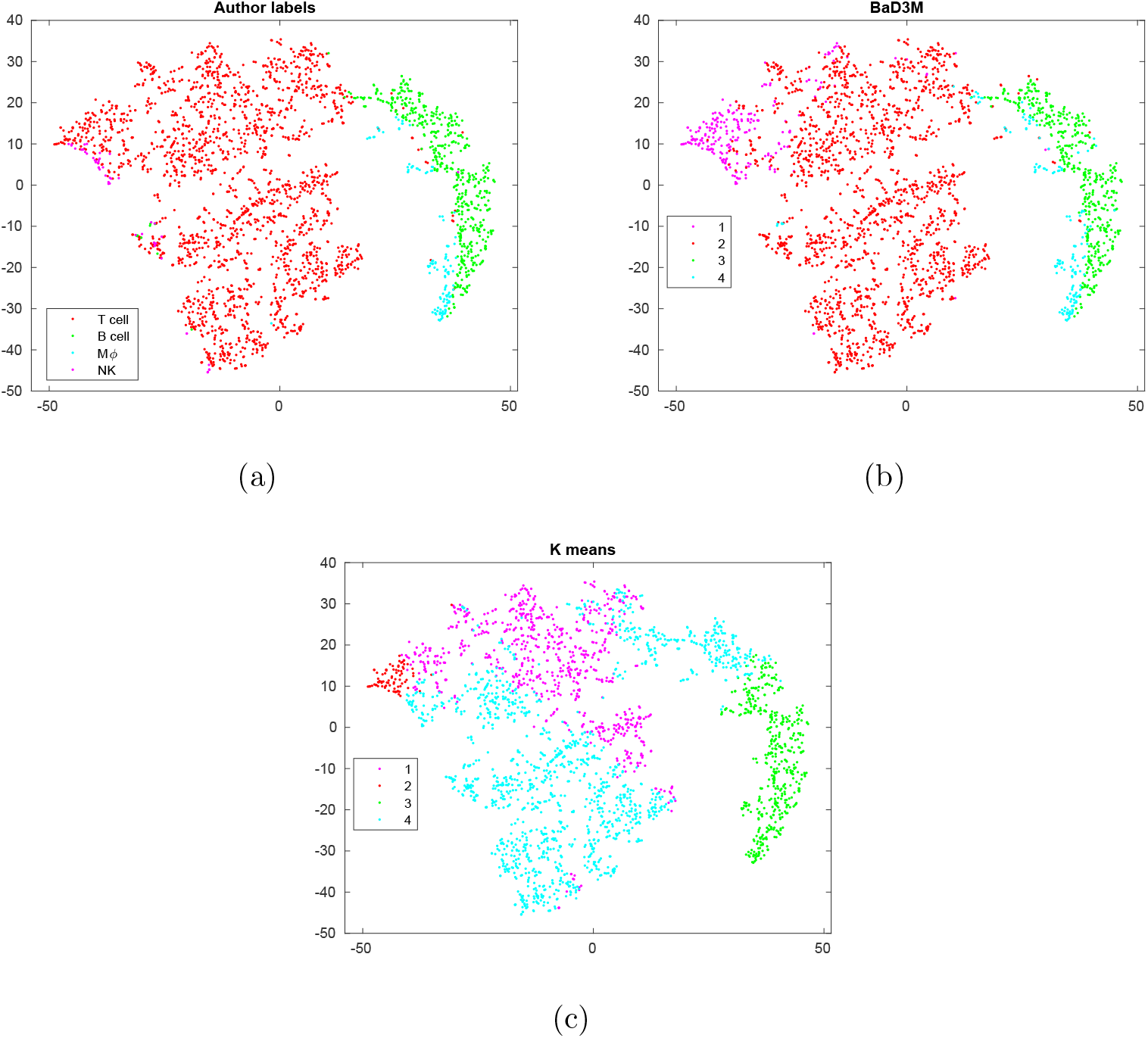
BaD3M automatically learns meaningful clusters. The cell type labels assigned by data collecting authors (a) and those inferred automatically through clustering with our proposed method (b) overlap well. The density based assumption behind k-means (c) fails to capture the nuanced differences among the various clusters. (M*ϕ*: macrophage, NK: natural killer)

The biggest region of disagreement is one group of T cells around around NK that our group co-clusters. To examine the results in more detail, we plotted the confusion matrices (Figure 8). Mostly, there is a strong degree of overlap between the clusters that our model learned and the cell type labels. However, there is one major group of 204 cells that the study authors called as T cells that our proposed method co-clustered with Natural Killer (NK) cells. This can be a mistake, or there can be some latent similarity that escaped the original author annotations, therefore we decided to investigate further.

**Figure 8:**
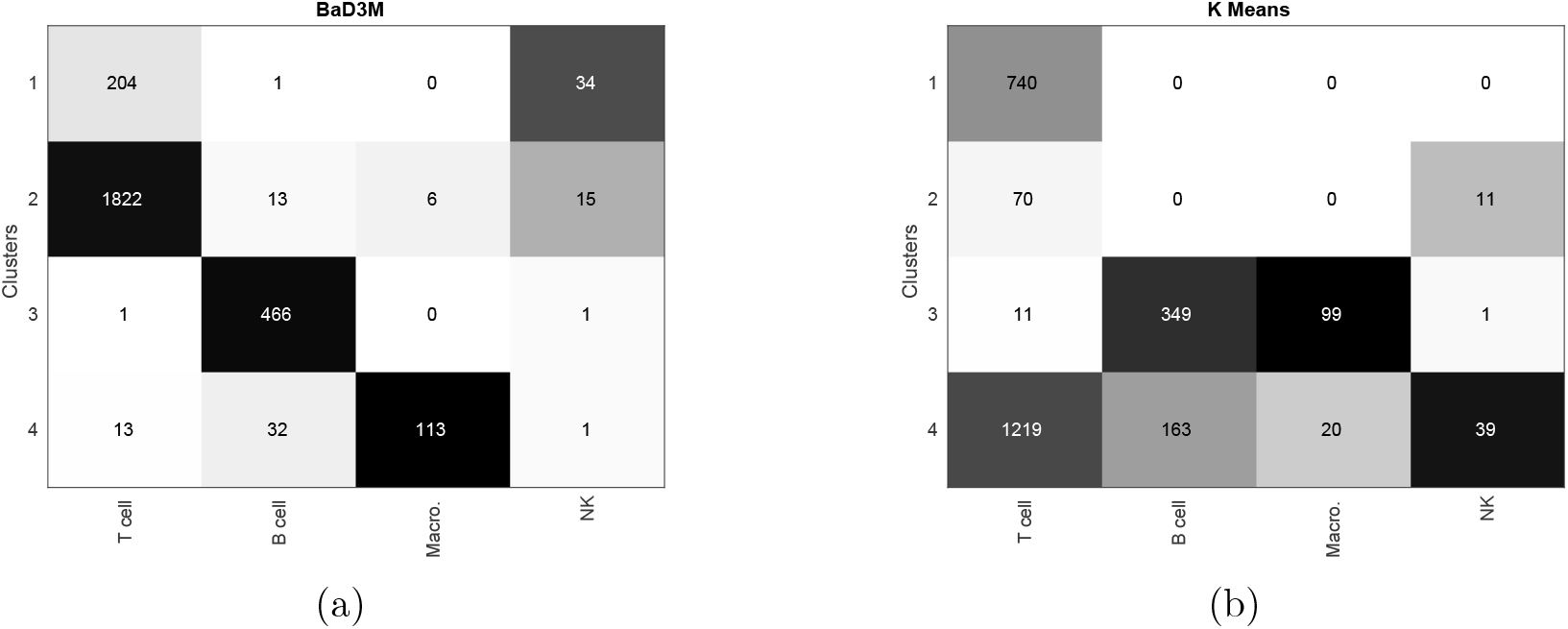
Confusion matrices of cluster assignments against author labels.

We used the probabilistic nature of our proposed method, which allows it to be interpretable, to plot the inferred gene expression tendencies of each cluster (formally, class-conditional Dirichlet-multinomial concentration parameters 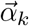) as shown in Figure 9. We can clearly see that the B-cell markers CD19 and MS4A1 (CD20) are correctly inferred to be high in a cluster of B cells, while CD14 is inferred as high in a cluster of macrophages. When we investigate cluster 1, which is the cluster with the most NK cell members, it is evident that CD8A is inferred to be heavily expressed for this cluster. This interpretability presents the intriguing possibility that these could be CD8^+^ NKT-like cells which share characteristics with both NK cells, CD8 T cells, and NKT cells (Wang *et al*., 2015). With k-means, there is simply no way to interpret the clustering results, which is generally the case with non-generative clustering methods.

**Figure 9:**
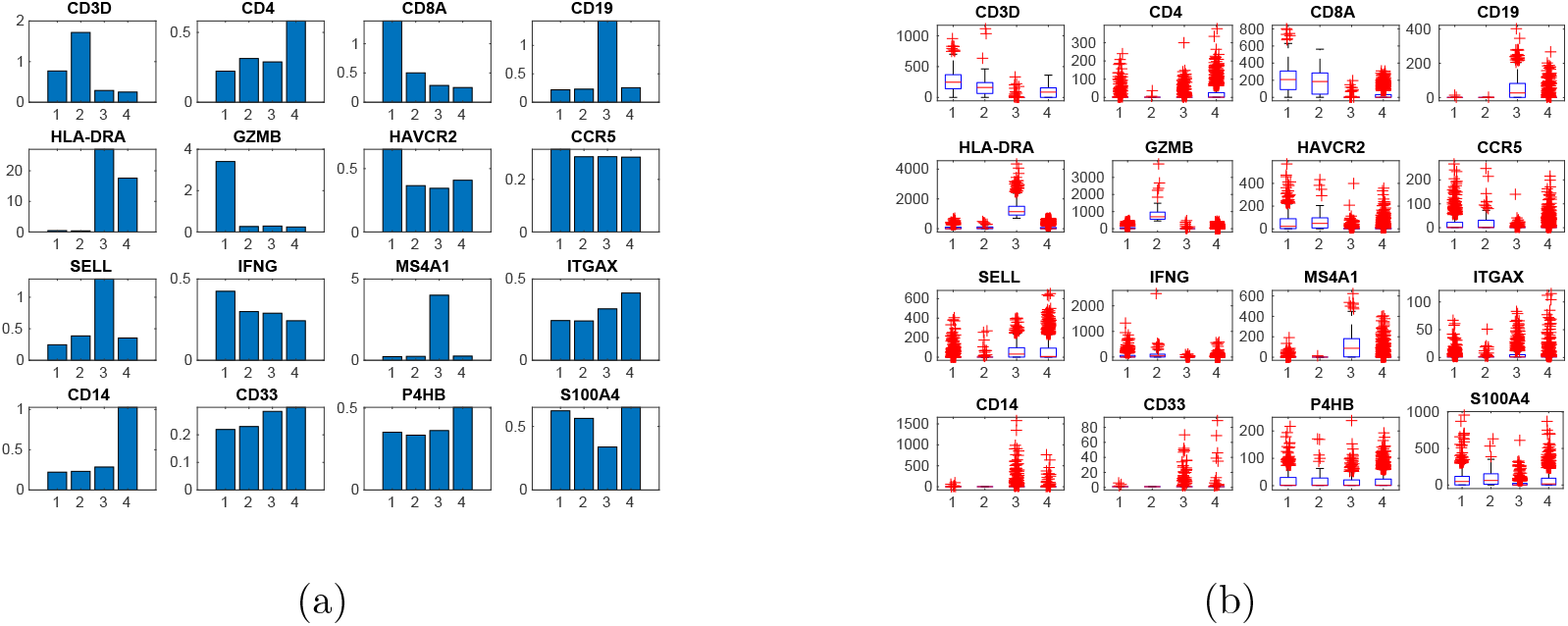
Marker genes in various clusters. (a) The expression tendencies of each marker gene for each cluster as learned by the proposed method. (b) The non-generative nature of K means does not allow us to interrogate the model, therefore we show the average expression of each marker in each cluster for K means.

## 4 Efficiency

Since one of our two key motivations was efficiency (along with interpretability), we would like to record the runtimes of the various models we discussed here. The runtime of our classification inference and predictions took under one second each for the 2,722 lymphocytes on a machine with 2.70GHz Intel Xeon E5-2680 CPUs (Figure 10). The automatic hyperparameter tuning ability of the SVM or GDA implementations led to their runtime being two orders of magnitude longer, otherwise these are also efficient algorithms. The inference is linear in its time complexity with respect to number of data points (provided a fixed number of iterations) as well as genes.

**Figure 10:**
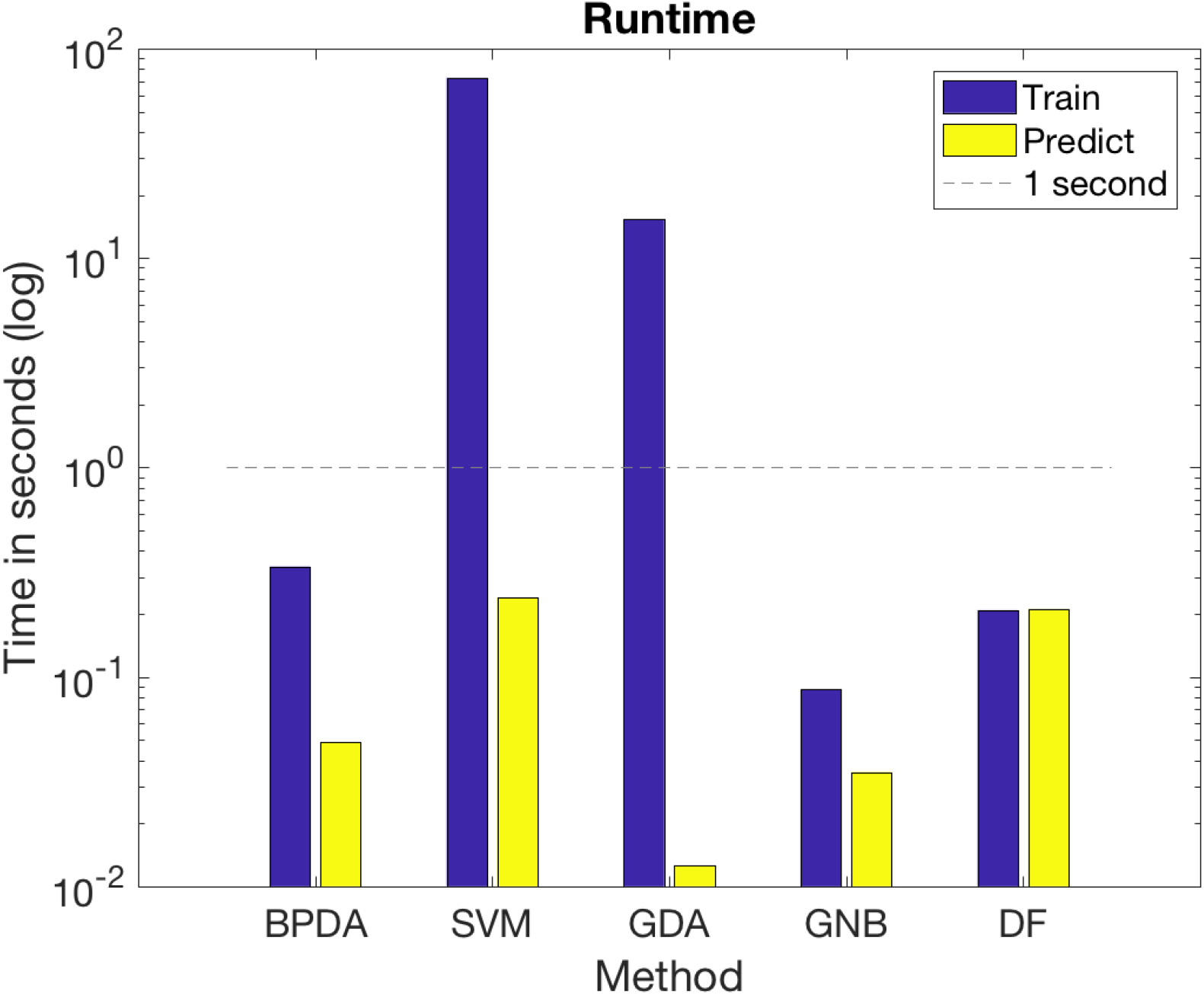
Run times of various classification methods.

In the clustering part, the algorithm took only 2.13 seconds on the same architecture mentioned above. In the E.M. algorithm, the E-step is assignment therefore it is linear time, hence fast. The maximization step could potentially be more problematic. However thanks to the optimization strategy we deployed (Newton-Raphson) the operations converge in a small number of steps compared to more popular approaches such as gradient descent. In addition the updates are also strongly optimized due to the Sherman-Morrison formula based trick we deployed, motivated by Minka (2012), which again renders time complexity linear instead of quadratic with respect to genes.

The number of iteration steps in the Newton-Raphson optimization is variable. In our implementation, we ran until either convergence, or a maximum number of iterations which we set at 40. In all the various trials we ran, Newton-Raphson always terminated due to convergence and never hit the iteration limit. Utilizing information from the Hessian in addition to the gradient enables NR to outperform the popular purely gradient based optimizations.

Thanks to the mathematical ideas we deploy here, these methods are highly scalable. The model’s memory requirements are *O*(*kg*) and *O*(*k*(*g* + 1)) respectively for the classification and clustering methods, where *k* is the number of cell types, *g* is the number of genes, therefore it is also scalable to store and share. Given all of the aforementioned arguments, we think that this method could scale to hundreds of millions of cells (such as the 300 million cells planned by the Human Cell Atlas) without difficulty.

## 5 Discussion

We present highly efficient and interpretable methods for two of the most central tasks in single cell RNA-seq analysis: namely cell type calling and cell clustering. Both methods share the same generative Dirichlet-multinomial assumption, yet solve two different problems, and therefore use different inference routines. The generative assumption renders our methods very easy to interrogate, as we demonstrate. Our proposed approaches are, to the best of our knowledge, the first fully probabilistic scRNA-seq analysis methods that are also highly scalable.

The ability of BDMC to cross the bulk/s.c. barrier means that it can enable the utilization of widespread purified cell population bulk RNA-seq data to enable efficiently labeling common cell types. The probabilistic nature of its predictions enables automated and information-theoretic (Shannon’s entropy based, for instance) analysis of these cell type calls. This would enable users to allocate more attention to finding and interpreting novel cell types.

The BaD3M clustering method also presents multiple novel advantages. To start with, the Dirichlet process class annotation means that it can be supplied with an upper limit of clusters (*K* = 100 or *K* = 30) and then be expected to set the mixture weights of these to zero, therefore automatically controlling cluster count. The mixture weights (i.e. 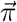) enable clusters with highly imbalanced memberships to be learned properly. Finally, the generative model enables making distinctions between various clusters that share many common genes but differ in the expression of only a few, but critical, genes.

In the future, we think that there are two main directions for improving the proposed methods. Firstly, the probabilistic assumptions can be modified to deal with the zero-inflated nature of single cell RNA-seq data. Zero expression renders multinomial based methods, which our methods are, numerically unstable and therefore we add a pseudocount of one to all expression values. As we show above, this does not cause severe performance drawbacks. However a more elegant solution would be the development of methods that natively represent the zero-inflated nature of this type of data.

Secondly, we think that the formalism we present can be extended to include pathway information. Specifically, it is possible to represent the expression profile of a cell as the juxtaposition of a number of shared pathways (metabolic, cytoskeletal etc) as well as cell type specific pathways (CD8 T cell specific, B cell specific etc). Our formalism easily extends to this by simply reparameterizing the class-onditional Dirichlet-multinomials with 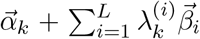 instead of simply 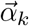 as we have now, where 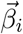 are one of *L* common pathways and 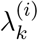 are the weights of each of these common pathways for a given cell type. The *α*_*k*_ in this model would learn exclusively class-specific expression programs, which would be even more informative if one is to interpret the learned models as a way of interpreting the novel clusters, for instance. These changes, while simple to state, imply significant inference challenges due to the coupling between *λ*, *α*, and *β*. Approximate inference methods such as sampling can be deployed, yet these would be multiple orders of magnitude slower than the methods proposed here. Therefore while such an extension would make the model more expressive and interpretable, it remains a major challenge.

In summary, we propose an efficient and interpretable approach for scRNA-seq data analysis. We hope that this provides researchers in the field with another tool at their disposal that they can deploy, especially as datasets with millions or hundreds of millions of cells are assembled. We hope that the collective analysis of such datasets can enable significant future discoveries in the life sciences.

## Acknowledgements

I would like to acknowledge useful discussions with Didem Ağaç Çobanoğlu who provided immunology domain expertise.

